# The brain compass: a perspective on how self-motion updates the head direction cell attractor

**DOI:** 10.1101/189464

**Authors:** Jean Laurens, Dora E. Angelaki

## Abstract

Head Direction cells form an internal compass that signals head azimuth orientation even in the absence of visual landmarks. It is well accepted that head direction properties are generated through a ring attractor that is updated using rotation self-motion cues. The properties and origin of this self-motion velocity drive remain, however, unknown. We propose a unified, quantitative framework whereby the attractor velocity input represents a multisensory self-motion estimate computed through an internal model that uses sensory prediction error based on vestibular, visual, and somatosensory cues to improve on-line motor drive. We show how context-dependent strength of recurrent connections within the attractor itself, rather than the self-motion input, explain differences in head direction cell firing between free foraging and restrained movements. We also summarize recent findings on how head tilt relative to gravity influences the azimuth coding of head direction cells, and explain why and how these effects reflect an updating self-motion velocity drive that is not purely egocentric. Finally, we highlight recent findings that the internal compass may be three-dimensional and hypothesize that the additional vertical degrees of freedom are defined based on global allocentric gravity cues.

## INTRODUCTION

Our ability to navigate through the environment is an essential cognitive function, which is subserved by specialized brain regions dedicated to processing spatial information (O’Keefe and Nadel, 1978; Mittelstaedt and Mittelstaedt, 1980). Of these spatial representations, head direction (HD) cells form an internal compass and are traditionally characterized by an increase in firing when the head faces a preferred direction (PD) in the horizontal plane (Taube et al., 1990a,b; Sharp et al., 2001a; Taube, 2007). In the rodent, the largest concentration of HD cells is found in the anterior thalamus (Taube, 1995; Taube and Burton, 1995), which supplies HD signals to other areas of the limbic system (Goodridge and Taube, 1997; Winter et al., 2015). The HD network is highly influenced by visual allocentric landmarks that serve as reference (Taube, 2007; Yoder et al., 2011), but is also able to memorize and update head direction in the absence of visual cues, e.g., when an animal explores an environment in darkness, by integrating rotation velocity signals over time. It is broadly accepted that these properties are generated through recurrent connections that form a neuronal attractor, which integrates rotation velocity signals (Redish et al., 1996; Skaggs et al., 1995; Zhang, 1996; Stringer et al., 2002). Although neuronal recordings support this concept (Peyrache et al., 2015; Kim et al., 2017), the underlying physiology remains poorly understood.

Lesions of the vestibular system disrupt HD responses, indicating that central vestibular networks are a major source of self-motion information (Stackman et al., 2002; Muir et al., 2009, Yoder and Taube, 2009; reviewed by Clark and Taube, 2012). However, neither the neuroanatomical pathways nor the computational principles of how vestibular cues influence HD cells have been worked out. Efforts to understand the functional principles of the links between the vestibular system and HD cells have been hindered by the lack of a theoretical perspective that unites well-defined (but traditionally segregated) principles about (1) how a neuronal attractor works, (2) basic vestibular computations (particularly those related to three-dimensional orientation) and visual-vestibular interactions, (3) how vestibular cues and efference motor copies are integrated during voluntary, self-generated head movements, and (4) recent findings on how HD networks encode head orientation in three dimensions (3D) (Calton and Taube, 2005; Finkelstein et al., 2015; Wilson et al., 2016). This review aims at presenting such a missing quantitative perspective, linking vestibular function relevant to spatial orientation with the properties of HD cells and the underlying attractor network. Our presentation is organized into four main sections.

First, we provide some intuition about the vestibular system’s role during actively-generated versus passive, unpredictable head movements, including a quantitative perspective of how rotational self-motion cues from vestibular, visual, somatosensory and efference motor copies interact (details can be found in Laurens and Angelaki, 2017). We then unite this self-motion model with a standard model of the head direction attractor in a single conceptual framework. We highlight that vestibular nuclei (VN) neurons with attenuated responsiveness during active head movements (reviewed in Cullen, 2012; 2014) carry sensory prediction error signals. Thus, they are inappropriate for driving the HD ring attractor, which explains why there are no direct projections from the VN to the lateral mammillary and dorsal tegmental nuclei (Biazoli et al., 2006; Clark et al., 2012), areas whose bi-directional connectivity is assumed to house the HD attractor (Basset et al., 2007; Clark and Taube, 2012; Sharp et al., 2001). Instead, we propose that the attractor velocity drive comes from the *final self-motion estimate*, which includes contributions of visual, vestibular, somatosensory and efference copy cues (Laurens and Angelaki, 2017). We discuss the implications of these properties for studying HD cells in freely moving or restrained animals, as well as in virtual reality. We highlight the role of the vestibular sensors and why, although the HD attractor drive is not exclusively driven by a sensory vestibular signal, vestibular lesions are detrimental to its directional properties.

Second, we summarize the fundamental properties of attractor networks, such that we can then disentangle the functional role of two factors: changes in the gain of self-motion velocity inputs to the attractor, and changes in the strength of recurrent connections within the attractor itself. We show through simulations that context-dependent changes on HD cell response properties, including experimentally reported decreases in HD cell modulation during passive/restrained versus active/unrestrained movements (Shinder and Taube, 2011; 2014a), can be reproduced through changes in the strength of recurrent connections within the HD attractor network. In contrast, changes in the gain of self-motion drive can explain the “bursty” activity, with drifting PD tuning, which has been reported previously in canal-lesioned animals (Muir et al., 2009).

Third, we summarize recent findings on how and why the animal’s orientation relative to gravity influences the properties of HD cells, including the loss or reversal of HD tuning in upside-down orientations (Finkelstein et al., 2015; Calton and Taube, 2005). We show that these experimental findings motivate a re-definition of the concept of azimuth during three-dimensional motion and further identify the three-dimensional properties of the self-motion signal that drives the HD attractor network. In fact, we show that egocentric velocity (e.g., yaw signals from the horizontal semicircular canals) alone cannot update the HD attractor. Instead, additional self-motion components encoding earth-horizontal azimuth velocity are necessary (Wilson et al. 2016).

Fourth, we highlight recent findings that HD cells may constitute a 3D (rather than azimuth-tuned only) compass (Finkelstein et al., 2015) and hypothesize that the additional vertical degrees of freedom may also have their primary drive from the vestibular system: in this case, by coding head orientation relative to gravity (Laurens et al., 2016).

Collectively, this review provides a quantitative framework that forms a working hypothesis on how self-motion cues update the HD attractor. Most importantly, we show that HD cell properties in both freely moving and restrained animals, as well as during planar exploration and 3D motion, are readily interpreted when these multiple elements, which are for the first time collectively presented together, are considered.

## NEURONAL ATTRACTOR

To begin, it is important to remember that there are three major components of the HD attractor (Redish et al., 1996; Skaggs et al., 1995; Zhang, 1996; Stringer et al., 2002; Peyrache et al., 2015; Kim et al., 2017), which are fundamental for understanding HD responses (Fig. 1A–C):

**Figure 1.**
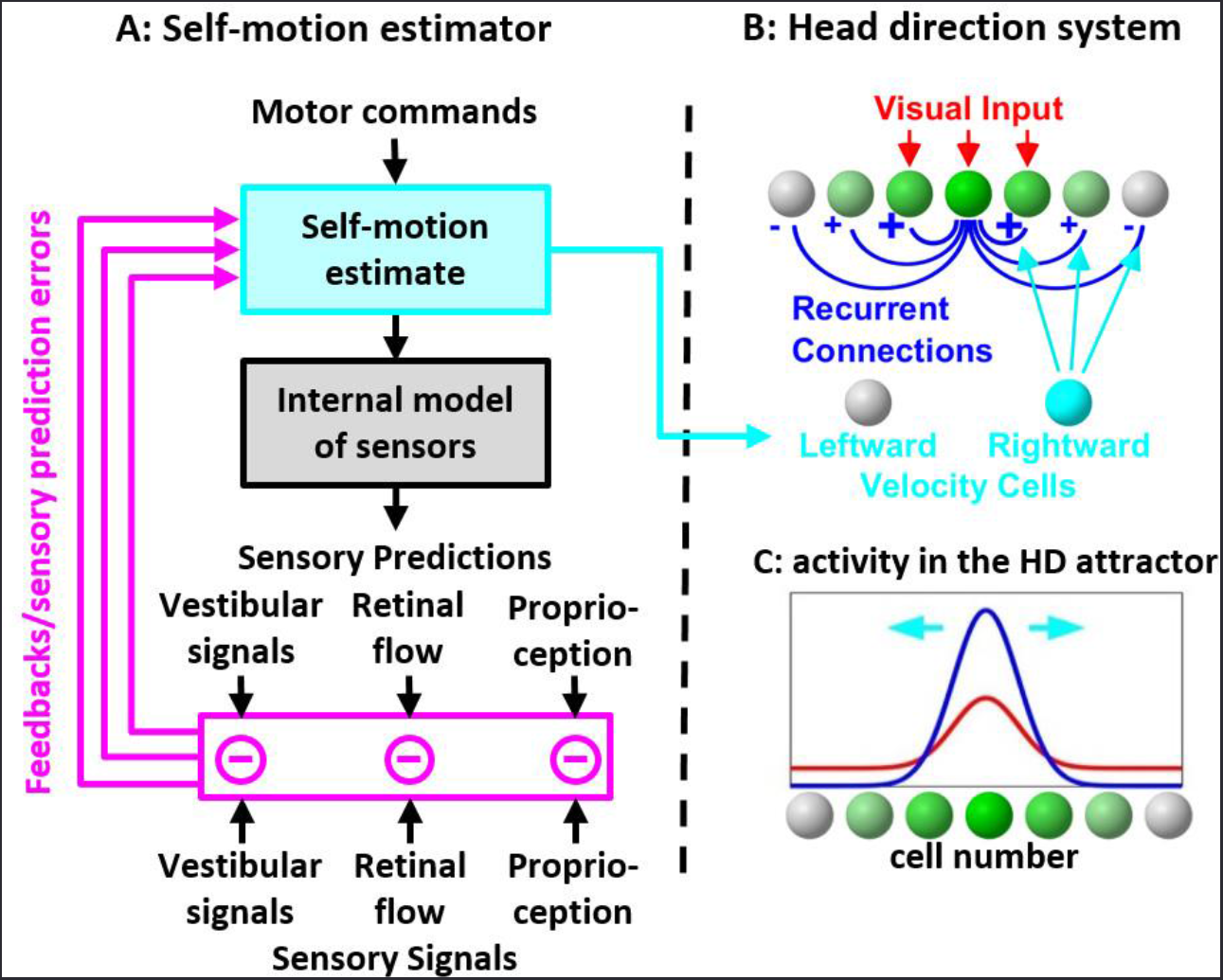
Self-motion drive to the head direction system. **(A, B)** Schematic of the self-motion estimator, where actual sensory signals (vestibular, proprioceptive, retinal flow) are compared with the corresponding predictions (generated through an internal model of the sensors) to generate sensory prediction errors (A, magenta) that improve the *self-motion estimate* (*cyan*). It is this *final self-motion estimate*, and neither the motor command nor the sensory prediction error (magenta), that has the appropriate properties (see simulations in Fig. 2) for updating the HD ring attractor (B). **(C)** Simulations of a HD ring attractor (Stringer et al., 2002) during random exploration. Activity of one neuron of the head direction attractor, under the influence of visual inputs alone (red) or with the additional influence of recurrent connections (blue). Self-motion signals result in a sideward shift of the hill of activity (cyan arrows).

1. A ‘landmark signal’ originating from the visual system activates cells whose PD corresponds to the animal’s current orientation in the environment (Fig. 1B).
2. Recurrent connections whereby neighboring cells activate each other and inhibit distant cells. Through mutual excitation, cells that are initially activated by the visual landmark signal activate each other even further, while inhibiting other cells. This mechanism amplifies the hill of activity and confers it the ability to sustain itself in the absence of visual inputs (Fig. 1C).
3. Self-motion velocity signals are thought to be conveyed by ‘velocity cells’ that influence the network by reinforcing recurrent connections in one direction, causing the hill of activity to slide in the corresponding direction (Fig. 1A,B, ‘self-motion signal’). This mechanism allows the attractor to integrate angular velocity over time.

## A STRAWMAN CONUNDRUM

We continue by discussing the properties of this self-motion velocity signal that provides the input to the HD cell attractor. In particular, we will address a confusion that exists in the field. As summarized above, lesions of the vestibular system disrupt HD responses, and head direction cell tuning disappears in vestibular deficient animals (Stackman et al., 2002; Muir et al., 2009, Yoder and Taube, 2009; reviewed by Clark and Taube, 2012), indicating that vestibular signals are required for the HD network to operate. On the other hand, actively-exploring animals generally have stronger HD tuning than passively-rotating, restrained animals (reviewed in Shinder and Taube 2014a,b), a property that has been interpreted as evidence that efference copies of motor commands are also a critical component of the velocity input to the HD ring attractor. Yet, normal HD tuning can be measured in restrained animals when they are rotated rapidly (Shinder and Taube 2011, 2014a), a fact that, again, suggests that vestibular signals play a primordial role in generating HD responses. Together, these observations create a puzzle that has never been clearly resolved. To make matters worse, there is a widespread misconception that vestibular signals are cancelled in actively moving animals. This notion has been created by inappropriately interpreting the functional implications of the fact that a particular (but prevalent) cell type in the VN shows reduced motion sensitivity during actively-generated rotations and translations (reviewed by Cullen 2012; 2014). This observation has led to the conundrum: if the vestibular pathway does not encode self-motion during active movements, how is the HD signal computed (Cullen, 2012; Cullen and Taube, 2017; Shinder and Taube, 2014b)?

Cullen and Taube (2017) offered two possible solutions to this conundrum: First, other cues, e.g., proprioceptive and/or neck motor signals that are available during active head motion, through multisensory integration, are ‘combined’ to provide a self-motion estimate. Although there is indeed no doubt that the velocity drive to the HD attractor represents a multisensory signal, how such multisensory integration can explain experimental findings has never been elaborated. In fact, in the classical framework for multisensory integration (Angelaki et al., 2009), the multisensory estimate is maintained by the remaining available cues in the absence of a sensory modality. But HD cell properties are destroyed without an active vestibular system, even in actively exploring animals. Thus, this phenomenological, qualitative solution does not give answers to the postulated conundrum.

The second explanation offered by Cullen and Taube (2017) is that the ascending signal to the anterior thalamus does not exclusively transmit angular head velocity, but also carries gaze-related signals. This hypothesis stems from the fact that neurons in the macaque nucleus prepositus hypoglossi, one of the pathways from the VN to the anterior thalamus, predominately carries eye-in-head position and velocity information (e.g., Dale and Cullen, 2013). Whereas the role of eye movements in the HD signal remains to be explored, this second explanation offered is also qualitative, without explaining why the contribution of gaze-relative signals can explain the fact that the absence of vestibular signals compromises HD tuning. Furthermore, neither of these solutions explain HD responses in restrained animals (Shinder and Taube 2011, 2014a).

Thus, in summary, both ‘solutions’ proposed by Cullen and Taube (2017) not only do not explain the experimental finding that HD cell tuning is compromised after vestibular lesions, but also highlight how inappropriate conclusions can be drawn when computational principles are not considered carefully when interpreting experimental findings. Instead, our position is that the conundrum described by Cullen and Taube (2017) (see also Shinder and Taube, 2014b) disappears when the interpretation that “vestibular self-motion signals are attenuated during active motion” is removed. In the only up-to-date modeling effort to understand how vestibular and extra-vestibular cues are combined during actively-generated head movements, we (Laurens and Angelaki, 2017) have proposed that the vestibular system is not only functioning, but also critically useful and important, for accurate self-motion sensation during both passive and active head movements. By considering a state of the art model of self-motion processing during active and passive motion, we are thus bridging a noticeable gap between the vestibular, motor control and navigation fields.

## INTERNAL MODELS, SENSORY PREDICTION AND ACTIVE VERSUS PASSIVE MOTION IN THE VESTIBULAR SYSTEM

Our insights come from a simple Kalman filter model, where we have extended a well-established framework developed previously for the optimal processing of passive vestibular signals using internal models (Laurens 2006; Laurens and Droulez, 2007; 2008; Laurens et al. 2010, 2011a; Laurens and Angelaki, 2011; Laurens et al. 2013a,b; Oman, 1982; 1989; Borah et al., 1988; Glasauer 1992; Merfeld, 1995; Glasauer and Merfeld 1997; Bos and Bles 2002; Zupan and Merfeld, 2002) to actively-generated head movements by incorporating motor commands (Fig. 1A; Laurens and Angelaki, 2017).

An important step in this model of multisensory self-motion estimation is the computation of sensory prediction errors (i.e., difference between actual and predicted sensory signals; Fig. 2B vs. 2C) that are used to correct the self-motion estimates. During active motion, motor commands can be used to predict head and body motion and anticipate the corresponding sensory re-afference, such that sensory prediction error is minimal (Fig 2D, top; Laurens and Angelaki, 2017). In contrast, sensory activity cannot be anticipated during passive/unpredictable motion, resulting in non-zero sensory prediction errors, which then drive the self-motion estimate (Fig. 2D, middle). We have shown that the simulated sensory prediction error mimics the properties of VN neurons with attenuated responsiveness during active motion (Cullen 2012; 2014). Importantly, even though sensory prediction errors differ, the final self-motion estimate is identical to head motion, irrespective of whether motion is active and passive (Fig. 2E), at least during short-lasting movements. Furthermore, if a passive motion (or a motor error) occurs during active movement (Fig. 2A-E, bottom), the vestibular organs will sense the total head motion (Fig. 2B, bottom). Within the central vestibular pathways, the internal model will predict the active component (Fig. 2C, bottom) whereas the VN neurons will encode specifically the passive component (or the motor error; Fig. 2D, bottom). The final self-motion signal (Fig. 2E, bottom) is the sum of the prediction (output of the internal model; Fig. 2C, bottom) and the error signal (Fig. 2D, bottom). The three simulations shown in the three rows of Fig. 2A-E also illustrate how the final motion estimate will ultimately be nearly identical (see Laurens and Angelaki 2017 for details) to the sensory signal (compare Fig. 2B and E) during short duration movements.

**Figure 2.**
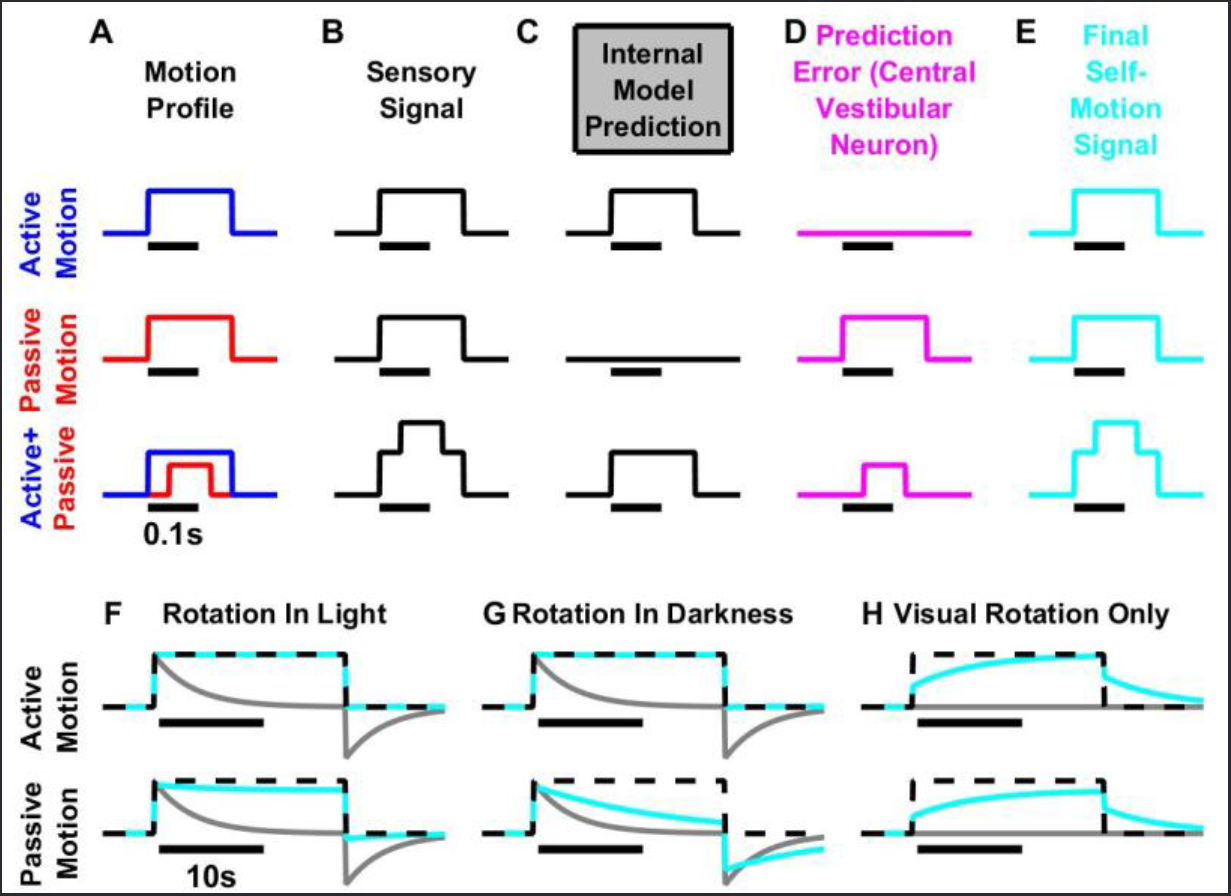
Internal model simulations during active and passive motion. **(A)** Two movements with identical velocity profiles (broken black lines) are performed actively (top), passively (middle) or a combination thereof (bottom). **(B-E) Fast head movements**. During active motion (top), the internal model uses the motor command to generate an accurate prediction of both self-motion velocity and sensory signal (C, top). The predicted sensory signal matches the actual signal, and therefore the sensory prediction error is null (D, top; magenta). During passive motion (middle), both the predicted self-motion velocity and sensory signal are null (C, middle), as there is no motor command. Therefore, the sensory prediction error is equal to the sensory signal (D, middle; magenta), which is transformed into a feedback that drives an accurate self-motion estimate. Thus, although the central vestibular neurons recorded by Cullen and colleagues, which encode sensory prediction error (Brooks et al., 2015), are largely suppressed during active motion, the *final self-motion estimate* is identical (and matches the stimulus) during both active and passive motion. For short rotations, simulation results are identical in light and darkness. Bottom line: Combination of active (blue) and passive (red) motion. The sensory signal (B) encodes the total (active+passive) motion. The internal model predicts the active component (C) and the sensory prediction error corresponds to the passive (D) component. The final self-motion estimate is identical to the total motion. **(F-H) Long-duration movements** (illustrated as constant velocity). Due to the mechanical properties of the vestibular sensors, the rotation signal (gray lines) decreases with a time constant of ~4s (in macaques) and exhibit an after-effect when rotation stops. The internal model of the sensors improves the self-motion estimate to a limited extent (Laurens and Angelaki, 2011; 2017). During rotation in light (F), visual rotation velocity cues create a sustained self-motion signal, thus the *final self-motion estimate* matches the stimulus during both active and passive motion. An accurate *final self-motion estimate* is also computed during active rotation in darkness (G, top). However, during passive rotation in darkness (G, bottom), the internal model cannot entirely compensate (see Laurens and Angelaki, 2011; 2017), and the *final self-motion estimate* decreases over time. **(H)** In visual virtual reality (VR; in this case, a rotating visual surround), the *final self-motion estimate* is inaccurate unless in steady-state (constant velocity) when the actual vestibular (canal) signal is null. Importantly, presenting the same visual stimulus while the animal attempts to rotate actively (but without simultaneous activation of the vestibular system) would induce a similarly inaccurate *final self-motion estimate* (G, top), even if other sensory cues (visual, somatosensory) and motor commands are available. These simulations highlight why the HD (and likely grid) signal is lost in VR. Simulations are based on the Kalman filter model of Laurens and Angelaki (2017).

The model of Laurens and Angelaki (2017) eliminates the misconception that vestibular signals are unimportant during self-generated, active head movements. The attenuated VN responses recorded by Cullen and colleagues (reviewed in Cullen, 2012; 2014) carry sensory prediction errors and are thus inappropriate to drive the HD attractor. In fact, there is currently no evidence to link VN response properties directly to HD cells, as there are no direct projections from the VN to the brain areas thought to house the HD attractor (Biazoli et al., 2006; Clark et al., 2012).

Instead, we propose that the *multisensory self-motion estimate* (Fig. 2D), rather than the sensory prediction error (Fig. 2C), updates the ring attractor (Fig. 1A, B). In fact, the model nicely recapitulates, for the first time, why vestibular cues are critical for HD tuning and why HD cells lose their spatial tuning properties in the absence of functioning vestibular signals, even during active head movements. This occurs because vestibular signals are continuously monitored to correct the internal model output and, without intact vestibular organs, the *self-motion estimate* which drives the HD attractor, would no longer be accurate during either active or passive motion. Thus, not only are vestibular sensory cues important, but also <u>absolutely necessary</u> for accurate self-motion computation during actively-generated movements. Simply because particular cell types do not modulate during active movements does not mean that the brain throws away vestibular signals during active motion. Sadly, this intuition has remained largely unappreciated, leading to the misunderstanding and the misguided strawman conundrum discussed by Cullen and Taube (2017).

The previous literature on HD cells has long recognized that proprioceptive and motor efference cues should participate, together with vestibular signals, to track head direction over time (e.g., see Clark and Taube, 2012: ‘internally generated information, or idiothetic cues (i.e. vestibular, proprioception, and motor efference), can be utilized to keep track of changes in directional heading over time’). However, a quantitative conceptual framework that describes the interplay between these sources of information has been lacking, and such statements have only been made in a qualitative manner. The model of Laurens and Angelaki (2017; simplified in Fig. 1A and 2) provides the missing theoretical framework of how multiple signals are combined to construct an appropriate multisensory self-motion estimate under all conditions (during active motion, as well as during passive motion, in light and in darkness) that then updates the HD cell attractor. Importantly, each cue’s contribution to the final self-motion estimate can be precisely predicted and quantitatively estimated based on the Bayesian framework.

Note that the final self-motion estimates are always identical during active and passive rotations in the light (Fig. 2E, F). However, during passive long-duration rotation in darkness (i.e., in the absence of motor commands, somatosensory and visual cues), the final self-motion estimate is inaccurate (Fig. 2G, bottom), because the vestibular sensors are poor at sensing low-frequency rotations. Accordingly, HD cells drift more during complex passive motion stimuli in darkness (Stackman et al., 2003). In contrast, during actively-generated movements, motor commands and/or available somatosensory cues improve rotation estimation and ensure HD stability even during long (several minutes) active foraging sessions in darkness.

Commonly-used rodent virtual reality (VR) environments, where the animal runs in place without the expected natural vestibular activation predicted by the internal model, lead to conflict conditions, where the vestibular sensory afferent signal does not match the sensory prediction. In this case, the *final self-motion velocity* (assumed to drive the HD ring attractor) is strongly underestimated during fast (high frequency) movements (Fig. 2H, early response). During slower or steady-state (low frequency) motion, however, retinal flow signals dominate self-motion perception and may mitigate the absence of vestibular signals (Fig. 2H, late response). This explains why HD and other spatial cell properties are altered in VR (Aghajan et al., 2015; Ravassard et al., 2013), as well as why results can be variable across studies and laboratories, depending on the visual VR stimuli and the running behavior of the animals.

In summary, understanding how the HD cell attractor is updated requires an understanding of the computational principles that govern self-motion perception and the fundamental role of sensory vestibular cues. Without the appropriate computational principles, it is easy to misinterpret experimental results. This misinterpretation has dominated the vestibular field in the past decade, and it would be a pity to propagate this misleading thinking into the spatial navigation forum too. Thus, we urge the spatial navigation researchers to rely on quantitative principles in order to understand how vestibular signals work together with other sensory and motor cues to generate (under most circumstances) accurate self-motion estimates that can update the multiple spatial cell representations in the navigation circuit. Although seemingly complex, there is a clear, quantitative, and logical framework that governs the roles of vestibular cues and their internal model for computing veridical self-motion estimates that are necessary to generate the spatial properties of HD, place and grid cells.

## RECURRENT CONNECTIVITY STRENGTH AMPLIFIES HD FIRING AND IS MODULATED BY BEHAVIORAL CONTEXT

Next we try to correct another incorrect attempt to link experimentally reported decreases in HD cell modulation during passive/restrained versus active/unrestrained movements and vestibular self-motion signals (Cullen and Taube, 2017). Shinder and Taube (2011; 2014a) recorded HD responses when (1) animals foraged freely, (2) animals were restrained (head and body) and rotated rapidly and continuously (>100°/s), (3) animals were restrained and placed statically (0°/s) in various orientations. In conditions (1) and (2), HD cells exhibited large firing rate modulation, firing in bursts (average 56 spk/s) when the animal faced the cell’s PD, and being almost silent (average 1 spk/s) when the animals faced away from it (Fig. 3A, blue and green curves). In condition (3), HD cells retained their PD, but their modulation range was reduced to 23 spk/s (in the PD) - 5 spk/s (away from the PD), (Fig. 3A, red symbols). Which property can account for and explain these experimental findings?

**Figure 3.**
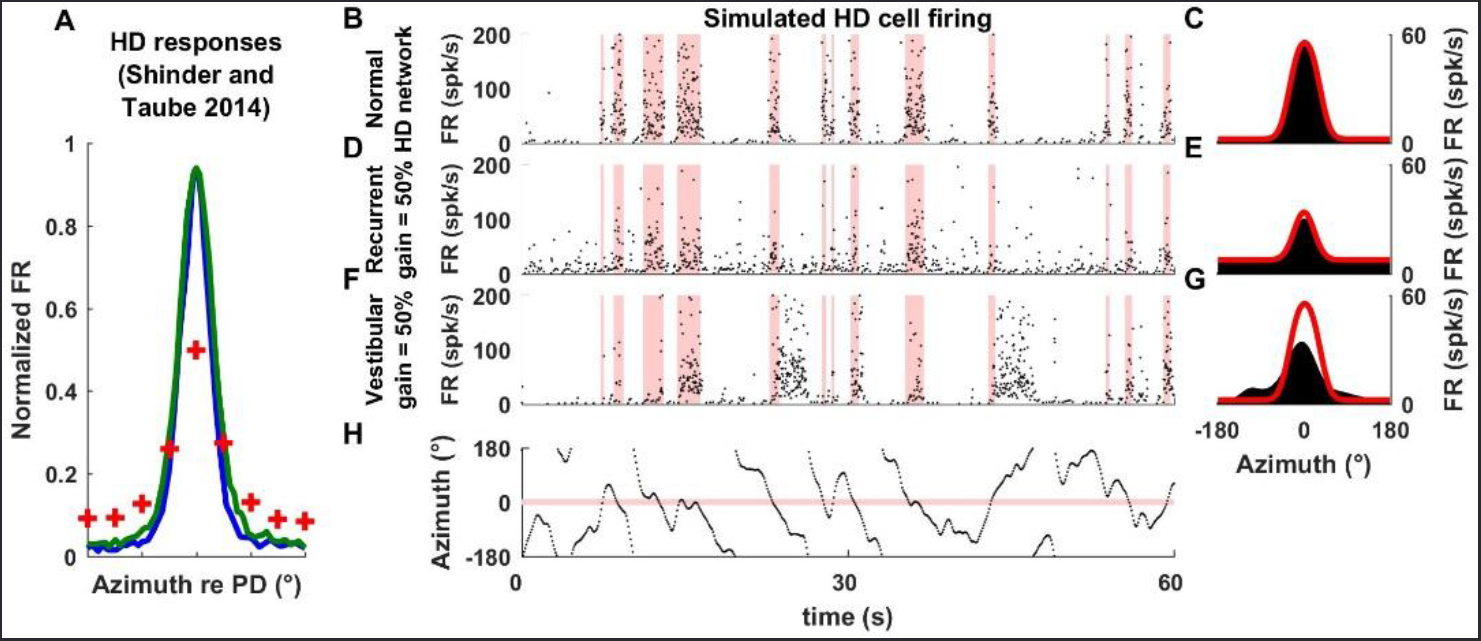
Context-dependent firing properties of HD cells. **(A)** Neuronal activity in HD cells reported in Shinder and Taube (2014a) when the animal moves freely (blue), is restrained and rotated rapidly (green) or restrained and placed at various static orientations (red crosses). **(B-H)** Simulated responses of a ring attractor cell under three contexts **(B,C)** when the unrestrained animal moves actively and freely, thus the attractor operates normally, **(D, E)** when recurrent connection strength is decreased by 50% (but self-motion velocity input remains the same), and **(F, G)** when the self-motion input gain that drives the ring attractor is decreased by 50% (but recurrent connection strength remains the same). Panels on the left illustrate simulated spiking activity (instantaneous firing rate). Pink bands indicate when head direction is within 30° of the cell’s PD. Panels on the right illustrate tuning curves reconstructed from the simulated firing on the left panels (black) or from simulations where the animal is placed statically in various orientations (red). Simulations are based on the model of Stringer et al. (2002). Note that when the animal moves freely, sharp bursts of firing occur when the head faces the cell’s PD, and the cell is nearly silent otherwise. When recurrent connection strength is reduced by 50% (D,E), the simulated model cell exhibits a higher background firing rate and lower modulation. When the gain of the self-motion velocity input signals is reduced by 50% (F,G), the activity in the HD attractor drifts relative to actual head direction. Thus, the cell’s burst of activity coincide only occasionally with the cell’s PD. **(H)** Motion profile used in the simulations in B,D,F.

Some investigators (Cullen and Taube, 2017; Shinder and Taube, 2014b) have considered that the attenuation of HD responses is related to the attenuation of neuronal responses in central vestibular pathways during active (as compared to passive) self-motion. However, several arguments can be raised against this interpretation.

1. As illustrated in Fig. 2 (and easily verified by behavioral studies), under most circumstances the final self-motion estimate, which is the signal that should update the ring attractor, is identical during active and passive motion.
2. Neuronal firing in the VN is attenuated during active motion (when HD cells respond consistently) and not during passive motion (where HD responses may be attenuated). Thus, the effects in the two cell types are opposite.
3. HD responses are attenuated in restrained animals, when placed statically in different directions, but are not attenuated in freely-moving animals that pause facing different directions. Yet, self-motion signals originating from the vestibular organs are equal to zero in both of these situations, regardless of whether or not the animal is restrained. Thus, this difference cannot be explained by considerations about the attenuation of VN responses during passive motion.
4. Indeed, notions of “passive motion” and “restrained animals” are often confounded. While all experiments on restrained animals use passive motion, passive motion can also be applied to freely moving, as well as, restrained animals. For example, Blair and Sharp (1996) rotated the platform onto which animals walked, and found that HD cells can track motion applied in this manner.
5. Finally, but most importantly, the restrained/passive motion findings of Shinder and Taube (2011; 2014a) are inconsistent with simulations based on an attenuated self-motion signal, as shown next.

Recall that recurrent connections within the ring attractor amplify HD responses by allowing cells with similar PDs to activate each other while inhibiting other cells (Fig. 1C). This property is further highlighted in Fig. 3B-G, which simulates both the instantaneous firing (left panels) and average azimuth tuning (right panels; black) of a HD cell. In a simulation where the strength of all recurrent connections is lowered by 50%, the HD cell retains its PD, but its peak firing rate is reduced and its baseline firing rate increases (compare Fig. 3D,E vs. 3B,C). Importantly, the cell exhibits the same response curve in a simulation where the animal is positioned statically for several seconds at each orientation (black vs. red curves).

In contrast, when the strength of recurrent connections remains unchanged but the self-motion velocity drive is “attenuated” by 50% (thus, inducing a mismatch between the velocity input to the ring attractor and the animal’s velocity), the packet of activity drifts relative to the actual head direction and coincides only occasionally with the cell’s PD (Fig. 3F, red vertical bars). The corresponding tuning (Fig. 3G, black) exhibits a weak peak at the cell’s PD (in this particular simulation, bursting occurs more often in the PD because visual information anchors the cell’s response to some extent). Although the average tuning curves in Fig. 3E and Fig. 3G (black) are somewhat similar, the instantaneous pattern of activity burst firing in Fig. 3D and Fig. 3F are entirely different.

We can now compare these simulations with the experimental findings of Shinder and Taube (2011; 2014a). The reduced responses observed in restrained/passive rotation experiments (Shinder and Taube 2014a), where HD cells respond consistently when the animal faces the PD but with a lower modulation amplitude, resemble the predictions of Fig. 3D (changes in the strength of recurrent connections), and not Fig. 3F (changes in the magnitude of input drive). Thus, we reason that restraining animals results in a decrease of recurrent connection strength in the HD attractor, probably though a tonic modulatory process. This recurrent activity may be restored by moving the animal rapidly (interestingly, even head translation may improve HD cell responses in the restrained animal; Shinder and Taube, 2011). However, the physiological mechanisms remain to be clarified.

Importantly, one should not conclude that vestibular signals are necessary to maintain recurrent connection strength. First, HD cells respond vigorously in unrestrained animals even when they stop moving for short periods of time (Shinder and Taube, 2014a; Taube, 1995; Taube et al., 1990a). Second, a study in sleeping rodents (Peyrache et al., 2015) indicates that activity in the HD network is identical during REM sleep and when animals are awake and freely-moving.

In parallel, the simulated firing pattern of Fig. 3F,G can explain the findings of Muir et al. (2009) and Yoder and Taube (2009) in animals with vestibular lesions. These studies reported “bursty” cells that alternate between periods of inactivity and short bursts of activity but bear little relationship to head direction. The alternation of inactivity and bursting resembles the activity of a neuronal attractor that would be decoupled from its velocity input and drift randomly (Clark and Taube, 2012), in line with the simulations in Fig. 3F. This activity can be further interpreted in light of the model in Fig. 1, which assumes that the *final self-motion signal* drives the HD attractor. We have reasoned that, although in principle this is a multisensory signal, vestibular lesions would lead to a mismatch with the sensory predictions during active head movements, thus destroy the normal correction feedback. As a result, the brain loses its ability to estimate self-motion accurately (accordingly, lesioned animals exhibit pronounced locomotor deficits), even though other sensory cues remain intact.

| <u>Vestibular Manipulation</u> | <u>HD findings</u> | <u>hypothesized effect</u> |
| --- | --- | --- |
| - Canal plugging<br>(Muir et al 2009) | bursty cells,<br>no HD tuning | ring attractor intact, self-motion estimate<br>compromised because sensory prediction<br>error is incorrect (Fig. 1A, 3E) |
| - Tilt mutant<br>(Yoder & Taube 2009) | bursty cells,<br>unstable HD tuning | ring attractor intact, self-motion estimate<br>mismatch (missing allocentric gravity<br>Component; see Fig. 6) |
| - Passive fast motion<br>(Blair & Sharp 1996;<br>Shider & Taube 2011;2014a) | normal HD tuning | ring attractor intact,<br>self-motion internal model intact |
| - Restrained animal<br>(Shinder & Taube 2014a) | reduced HD tuning | reduced recurrent collaterals of ring<br>attractor, self-motion estimate intact |

In summary, the HD activity attenuation observed in some studies is most likely due to the restraining (not the self-motion signal itself) altering the strength of recurrent connections of the ring attractor. Furthermore, collectively the simulations of Fig. 1–3 illustrate that the issue of activity attenuation in some central vestibular neurons during active and passive motion (Cullen, 2012; 2014) is completely irrelevant and largely disconnected from that of the factors that contribute to HD cell activity during active vs. passive motion, or in restrained vs. freely moving animals. Making such links only side tracts and mis-directs our attempts to understand how self-motion influences the spatial properties of HD (and grid) cells.

Having, as we hope, corrected the mis-representations of what the important issues are in linking vestibular and, more generally, multisensory self-motion signals to HD tuning, we next highlight the true challenges regarding the properties of the self-motion drive of the HD attractor. We next show that this multisensory self-motion velocity input to the HD attractor has more complex spatial properties that originally envisioned.

## GENERALIZED DEFINITION OF AZIMUTH COMPASS THAT MAINTAINS ALLOCENTRIC ORIENTATION DURING THREE-DIMENSIONAL MOVEMENT

Recall that HD cells have been traditionally recorded during motion in an earth-horizontal plane. As such, the issue of reference frames was trivial. For example, self-motion cues updating the ring attractor could be purely egocentric; e.g., yaw velocity signals from the horizontal semicircular canals (Calton and Taube, 2005; Taube, 2007), as long as visual landmark cues anchor HD tuning to the allocentric local environment (Taube, 2007). Importantly, when limited to the horizontal plane, the local and global (the latter defined as the earth’s surface) allocentric frames coincide.

More recently, a few studies have allowed animals to move in 3D (Calton and Taube, 2005; Finkelstein et al., 2015; Wilson et al., 2016), thus forcing the question of whether self-motion velocity signals that update the HD attractor are represented in egocentric or allocentic reference frames (Taube and Shinder, 2013). But before we summarize experimental findings, one must first appreciate that neither solution is without problems.

Let’s consider an allocentric azimuth updating rule first (Fig. 4A). One way to define azimuth during 3D head motion could be to project the direction the head faces (red arrow) onto a compass in the earth-horizontal plane. However, this solution distorts the angles between points in the head-horizontal plane. This is illustrated in Fig. 4B, where the diagram in A is viewed from the top. Red, blue and green arrows represent 0°, 90° and 45° leftward directions on the head-horizontal plane. When projected onto the earth-horizontal plane, the red and blue arrow will point North and West, i.e. 90° apart, but the green arrow will point between West and North-West, i.e. 71° from the red and 29° from the blue arrow. Thus, defining azimuth by projecting directions onto a earth-horizontal plane induces a distortion that would severely complicate the ability to orient oneself when navigating on a sloped surface, because the orientation of various landmarks in a visual reference frame, or the amplitude of head rotations in the sloped surface’s plane, would not correspond to the azimuth signal stored in the HD network. Furthermore, earth-vertical surfaces, like walls of a box, would not be represented at all.

**Figure 4.**
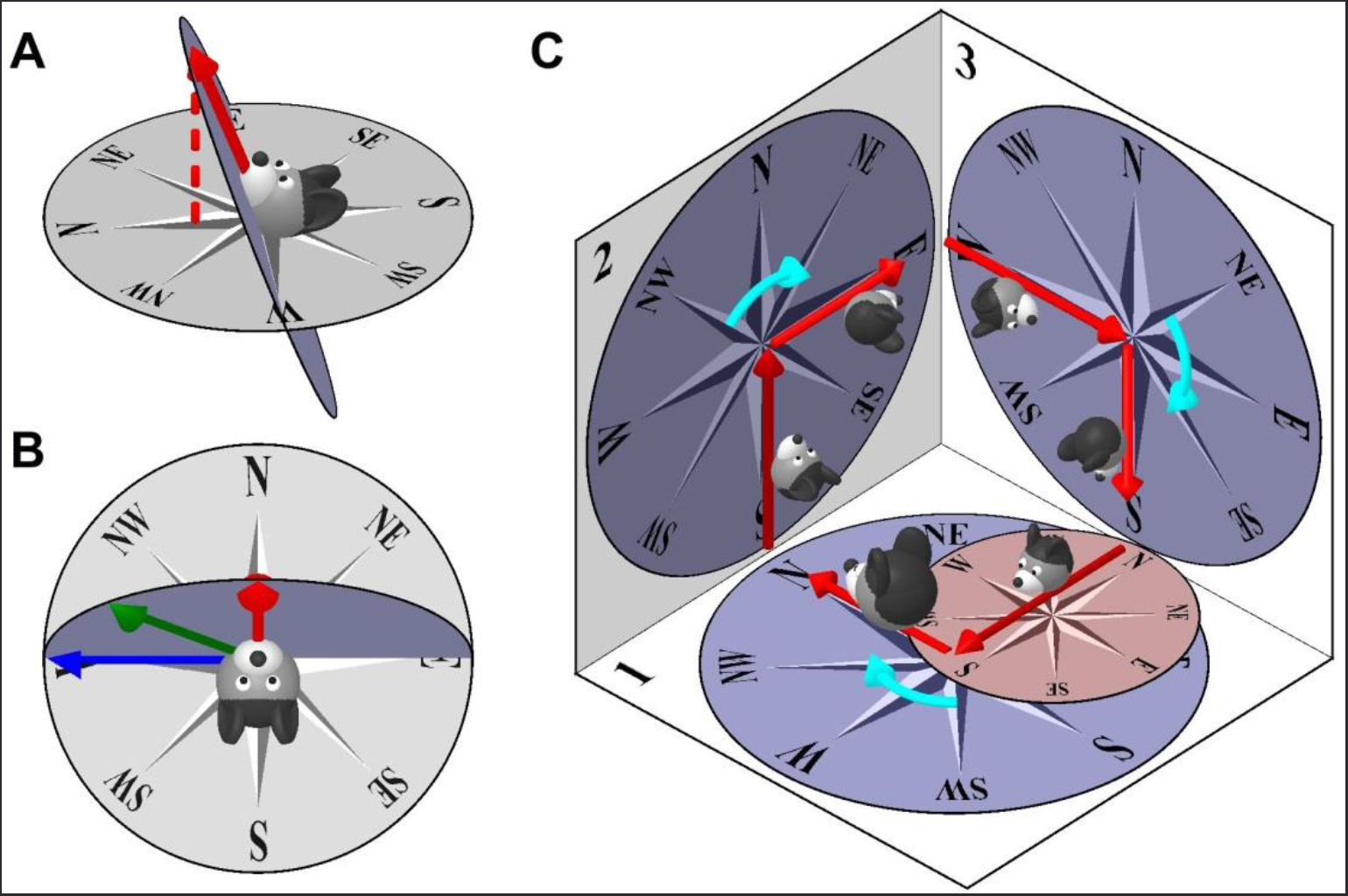
Earth-horizontal (Allocentric updating) vs. Head-horizontal (Egocentric updating) azimuth models. **(A), (B)** Earth-horizontal azimuth model, illustrating the distortion produced. The larger the tilt angle of the head away from the earth-horizontal surface, the larger the distortion between egocentric and allocentric reference frames. **(C)** Head-horizontal (yaw-only) azimuth model, with the compass attached to the head horizontal plane. After completing a 3D trajectory and returning to surface n°1, the head-fixed compass (small red compass on surface n°1) is no longer consistent with its orientation at the beginning of the trajectory (large compass on surface n°1). Therefore, a purely egocentric azimuth compass can’t track head orientation during motion in 3D.

One may then hypothesize that the solution is to use an egocentric azimuth updating rule (Calton and Taube, 2005; Taube and Shinder, 2013) where azimuth would be updated based on egocentric yaw rotation signals only, and remain unchanged during pitch and roll movements. Unfortunately, as illustrated in Fig. 4C, this solution would also fail, this time even more seriously. In this hypothesized motion trajectory, the rodent is initially facing North on an earth-horizontal surface (n°1 in Fig. 4C, large head) and then moves across two earth-vertical surfaces (n°2 and n°3) before coming back to its initial orientation. When an animal initially faces North on an earth-horizontal surface (n°1) and pitches nose-up, the compass would simply follow the rotation of the head and the North axis would now point upward. This solution would allow the animal to navigate naturally on the vertical surface. Then the animal turns East to proceed along surface n°3 and pitches 90° when it enters it. At this point, he faces East (since azimuth is not updated when he pitches from surface n°2 to n°3), and the compass on surface n°3 is drawn according to this orientation. After some time, the animal turns again and faces South, goes through another pitch of 90° in order to return to surface n°1. At this point, the internal azimuth signal (small red compass on surface n°1) would incorrectly encode a Southward orientation, although the animal is in fact facing West. When the animal returns to its initial orientation (large head on surface n°1), it will have performed exactly three 90° rightward horizontal turns (cyan arrows, when on each surface), interleaved with another three 90° nose-up pitch turns (when it transitions from one surface to another). A purely egocentric azimuth model (updated by horizontal canal signals only) would update during the horizontal turns, such that the total rotation registered by the HD attractor would be 3x90=270°. However, the correct attractor should have registered 360°. What went wrong in the arithmetic? This example illustrates that a purely egocentric azimuth updating rule (using yaw velocity signals only) cannot track head orientation during 3D motion.

Given that both egocentric and allocentric azimuth updating rules face serious limitations (Fig. 4), which solution is used by the brain? We summarize the experimental findings next (Fig. 5).

**Figure 5.**
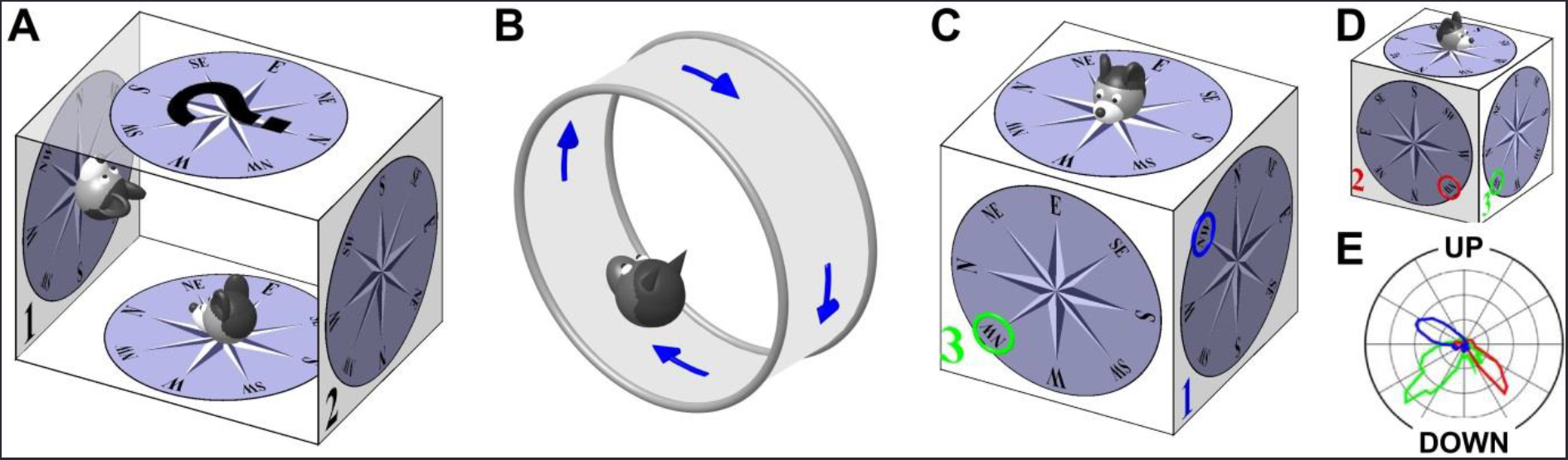
Summary of experimental findings on HD responses during locomotion on surfaces that are not earth-horizontal. **(A)** Experiment by Calton and Taube (2005). Rats are placed inside a box and walk on the floor, two vertical walls (n°1 and n°2) and ceiling. HD tuning is maintained and anchored to the head-horizontal compass; except when walking on the ceiling. **(B)** Experiment by Finkelstein et al. (2015). Bats crawl along a vertically oriented circular track. **(C), (D)** Experiment by Wilson et al. (2016), where rodents walk on the outside surface of a cube. Three surfaces are numbered (n°1, 2, 3) and color-coded. The NW direction is indicated by a colored circle on each surface. (D shows the same cube as in C, seen from another angle to display side n°2.) **(E)** Polar representation of the tuning curves of an example HD cell on sides n° 1-3 (color-coded), as seen from a camera outside the cube. Note that the cell’s PD on various sides corresponds to different directions in space. However, these directions correspond to the same azimuth (NW) in the tilted reference frame defined in Fig. 6.

1. Calton and Taube (2005; see also Stackman et al., 2000) trained rats to walk along the inner surface of a box. Starting from the bottom surface, they walked up one wall (marked n°1 in Fig. 5A), then walked upside down across the ceiling, climbed down the opposite wall (marked n°2), and reached the bottom floor again to receive a reward. HD cells in the anterior thalamus, which were first characterized on the bottom surface of the box, also responded when the animals walked on both vertical walls. When an animal walked facing North onto wall n°1, its orientation continued to be coded as North. Furthermore, when the animal descended wall n°2, its orientation still coded North. The authors concluded that HD cells encoded orientation as if the entire surface of the bottom and walls were a single continuous and flat surface (’locomotion plane’). Therefore, these findings are consistent with the egocentric updating rule (e.g., from the horizontal semicircular canals). When animals walked upside-down on the ceiling, HD tuning was either lost or bore no relation with the HD tuning recorded on the floor.
2. Finkelstein et al. (2015) recorded HD cells in the dorsal pre-subiculum of bats that crawled around the inner surface of a vertically oriented circular track (Fig. 5B). In agreement with Calton and Taube (2005)’s hypothesis that the azimuth reference frame extends to the whole locomotion surface, they found that, if the animal faced a cell’s PD on the bottom of the track, the cell would exhibit a large response during the entire motion along the track, including at the top. Thus, when walking upside-down, some HD cells appear to reverse PD (since, in an allocentric frame of reference, the animal faces opposite directions on the floor and on the ceiling), whereas other cells lost their tuning. The finding that azimuth tuning reverses in inverted animals persisted when animals were wrapped in a towel and displaced manually upside-down. Thus, as with the previous findings of Calton and Taube (2005), these results are consistent with an egocentric azimuth updating rule.
3. Finally, a major breakthrough came with the experiments of Wilson et al. (2016), who trained mice to walk on top and along all four vertical walls of a cube (Fig. 5C–E). When neuronal responses were recorded on opposite walls of the cube, PDs reversed (in allocentric coordinates), as in previous experiments. When animals transited between adjacent sides of the cube (e.g. from surface n°2 to n°3 in Fig. 5C), PD changed by 90° (Fig. 5E), in contrast to the predictions of the purely egocentric updating rule, which would predict no change. Why does this happen and what do these findings suggest regarding the updating rule for the HD ring attractor?

Recall that an egocentric azimuth solution fails because it loses its allocentric invariance when moving outside the earth-horizontal plane (Fig. 4C). Next we describe the solution, which we refer to as the *“tilted azimuth compass”*, which ensures that the head-azimuth compass can always update its allocentric orientation during 3D motion.

We reason that the correct (*tilted azimuth*) compass must be updated when animals transition from one earth-vertical surface to another (i.e., *when the head-horizontal plane rotates around an earth-vertical axis*). In Fig. 6A (replotted from Fig. 4C), for example, this includes the transition from surfaces n°2 to n°3 (but not the transitions from surfaces n°1 to n°2 or n°3 to n°1). This is because, although the animal is facing East on surface n°2, it faces South when it enters surface n°3. With this 90° correction (which clearly HD cells register; Fig. 5E), the ring attractor would correctly update by 360° in the trajectory of Fig. 6A. Therefore, the correct input drive to the azimuth ring attractor should include two components (Wilson et al., 2016): (1) a *head-horizontal* (yaw) rotation velocity (i.e., rotation in the head-horizontal plane), as originally assumed; and (2) an *earth-horizontal* rotation velocity (i.e., rotation around an earth-vertical axis; Fig. 6A, broken green line and green arrow).

**Figure 6.**
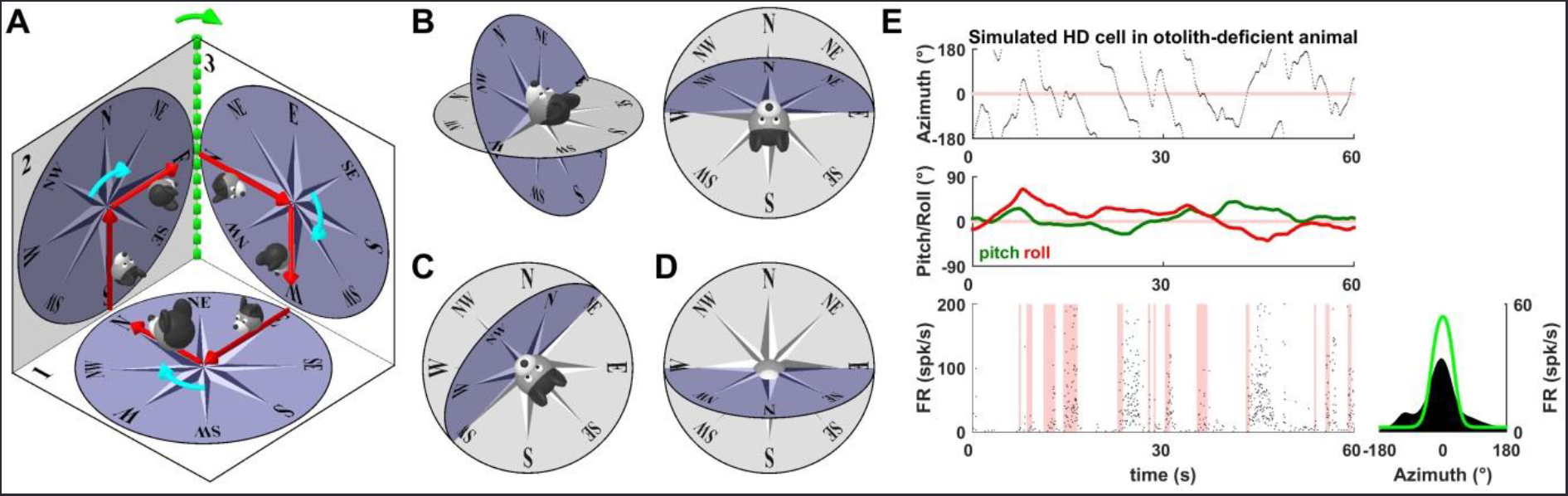
Definition of tilted azimuth model. **(A)** Consistency of the tilted azimuth compass with both egocentric and allocentric reference frames during 3D motion (multiple surfaces). Starting from a north orientation on an earth-horizontal surface (initial position marked by a large head), the animal follows a trajectory and comes back to its initial orientation (as in Fig. 4C). The trajectory includes three right-hand turns (in the head-horizontal plane, cyan arrows), each of which changes head azimuth by 90°. Azimuth must also be changed by 90° when the animal moves from surface n°2 to n°3 (rotation about an earth-horizontal axis, green) to ensure a 360° total azimuth change when the animal returns to its initial orientation. **(B-D)** Comparison of the tilted azimuth model (blue; Wilson et al., 2016) and the earth-horizontal frame (gray). **(B)** Example orientation, where the N, S, E and W directions of the head-horizontal and earth-horizontal compasses match, but not the NE and NW (as well as SE and SW). **(C)** The head-horizontal plane now intersects the earth-horizontal plane along the NE-SW axis. Accordingly, the two compasses now match along these directions, as well as in the NW direction, but not along the N and S directions. **(D)** Illustration of the tilt angle exceeding 90°, where the azimuth reverses along the N-S axis: the N direction of the head-horizontal compass is now aligned with the earth-horizontal S direction. **(E)** Simulation of a HD cell (as in Fig. 3) when the animal tilts its head (pitch/roll) assuming that the rotation drive signal to the HD ring attractor doesn’t account for head tilt (i.e., head-horizontal azimuth compass, updated only by yaw rotations). Cell firing doesn’t correspond to the cell’s PD consistently, resulting in drifts, as reported in otolith-deficient mice (Yoder and Taube, 2009). Light green: azimuth tuning curve obtained if head tilt is accurately accounted for (as is the case with the tilted azimuth model).

This solution (equivalent to the solution proposed by Wilson et al., 2016) ensures that azimuth on the earth-horizontal and head-horizontal compasses always remain anchored at the level of the line where the two planes intersect (i.e. E-W axis in Fig. 6B, NE-SW axis in Fig. 6C). The compromise, however, is that increasing tilt angle beyond 90° leads to a reversal of azimuth, i.e. the North direction in the head-horizontal compass (Fig. 6D, blue) is now aligned with the South direction in the earth-horizontal compass (gray). This explains the reversal of azimuth reported by Finkelstein et al. (2015) and Calton and Taube (2005). This reversal occurs because, in Fig. 6D, the East and West directions are anchored to the earth-horizontal East and West directions. In contrast, the North and South directions in the tilted frame are not earth-horizontal. As a result, instead of being anchored to earth-horizontal directions, they are defined relative to East and West. By definition, North lies 90° CW relative to West and 90° CCW relative to East in the plane of the compass. Placing it in the corresponding location in Fig. 6D results in North pointing in a Southward direction in the earth-horizontal plane.

Importantly, to update the azimuth ring attractor properly when the head-horizontal plane changes in space, the velocity drive to the HD circuit must include spatially-transformed, allocentrically (gravity)-defined self-motion signals. Gravity-referenced signals are indeed found in both the VN and cerebellum (Angelaki et al., 2004; Yakusheva et al., 2007; Laurens et al., 2013a,b).

These same principles apply when an animal walks on a horizontal surface but tilts its head. To illustrate the importance of this transformation, we simulated the same cell as in Fig. 3 but assumed that the head was tilting randomly (Fig. 6E, top) and the brain was not compensating for this tilt, but instead the HD attractor was updated by only egocentric rotation signals in the head-horizontal plane. Similar to the cell in Fig. 3E, the cell in Fig. 6E fires in bursts of activity that are inconsistently aligned with the cell’s PD. This simulation may explain findings in otolith-deficient mice (Yoder and Taube, 2009) where ADN neurons exhibited the pattern of “bursty” activity described in Fig. 6E, indicating that the HD signal was unstable. We propose that the lack of functioning otolith organs in these animals prevented the spatial transformation of rotation signals and the computation of the allocentric velocity component. This would induce a mismatch between the velocity input to the ring attractor and the animal’s velocity, resulting in activity bursts that drift relative to actual head direction, exactly as demonstrated experimentally by Yoder and Taube (2009). The tilted azimuth framework also accounts for the observation that the direction-specific discharge of HD cells was usually not maintained when the rat locomoted on the vertical wall or ceiling in 0-G (parabolic flight; Taube et al., 2004).

Thus, in summary, the velocity self-motion signal updating the azimuth compass (i.e., the drive to the HD ring attractor) is more complex than originally envisioned. Because of the need for continuous matching between head-horizontal and earth-horizontal (allocentric) compasses, the animal’s orientation relative to gravity must be monitored and used to transform self-motion rotation signals. Vestibular signals are critical for this transformation too.

Note that, in Fig. 6A, the azimuth is North if the head faces upward on surface n°2 but East if it faces upward on surface n°3 – which seems paradoxical since the head is facing the same allocentric direction. This is because the titled azimuth does not encode “upward” or “downward” directions, nor does it encode head tilt (although tilt signals are required to update it). Thus, in addition to the *tilted azimuth compass*, which needs gravity signals, the HD system may depend on gravity even more, as it may monitor the animal’s 3D orientation in the world. Next we summarize what little is currently known about the 3D properties of HD cells.

## IS THERE A THREE-DIMENSIONAL COMPASS – AND DOES IT DEPEND ON GRAVITY?

Can the HD system represent all three dimensions of head orientation in space, or is it just an one-dimensional (tilted) azimuth compass that can maintain its allocentric reference? For example, in Fig. 6A, the animal faces North when it looks upward on surface n°2, and East if it looks upward on surface n°3. If a 3D compass exists, then some HD cells should fire preferentially when the animal faces upward, independently of azimuth.

Indeed, a recent study in bats showed that HD cells in the pre-subiculum are tuned in 3D. Specifically, some cells signal the angular orientation of the head in pitch and roll (i.e. 2D head tilt), independently of azimuth, and some cells carry both tilt and azimuth signals (Finkelstein et al., 2015). However, Finkelstein et al (2015) did not consider reference frames (Fig. 4, 6) and mathematical issues related to computing 3D orientation and instead restricted their analysis of tilt and azimuth coding by excluding tilt movements to the side (e.g. roll).

Note that encoding 3D head orientation in a 3D attractor would raise a “combinatory explosion” issue (because of the large number of cells required to represent 3D orientations, and the large number of connections required to encode all possible rotations from one orientation to another). One solution to avoid this mathematical complexity, is to encode head azimuth independently from the two other degrees of freedom, and use gravity to define these remaining degree of freedom, i.e. vertical head tilt orientation. In fact, since correctly updating the azimuth HD ring attractor requires knowledge of orientation relative to gravity (Fig. 6), this solution does not require much additional computation.

Indeed, Laurens et al. (2016) have recently described pitch- and roll- coding cells that were anchored to gravity, and not visual cues, in the macaque anterior thalamus. Preliminary findings show gravity-tuned cells in the anterior dorsal thalamus of mice, with many cells being jointly tuned to both azimuth and gravity (Cham et al. in preparation), exactly as previously described in the bat presubicculum (Finkelstein et al., 2015).

Thus, as outlined in Fig. 7A, we propose that the HD system may form a 3D compass, where two dimensions are defined by gravity (2D tilt, spherical topology, green) and the other dimension by the tilted azimuth compass (circular topology implemented by the ring attractor, blue). Importantly, gravity signals influence computations for both the 2D tilt and tilted azimuth updating. In fact, the same internal model computations for self-motion estimation shown in Fig. 1 can also simultaneously compute tilt and gravity signals (Laurens et al., 2013a,b; Laurens and Angelaki, 2017). The vestibular system on its own can sense head tilt unequivocally - but not head azimuth. This explains why visual signals are necessary to anchor azimuth signals, but are not necessary for tilt signals (Laurens et al., 2016).

**Figure 7.**
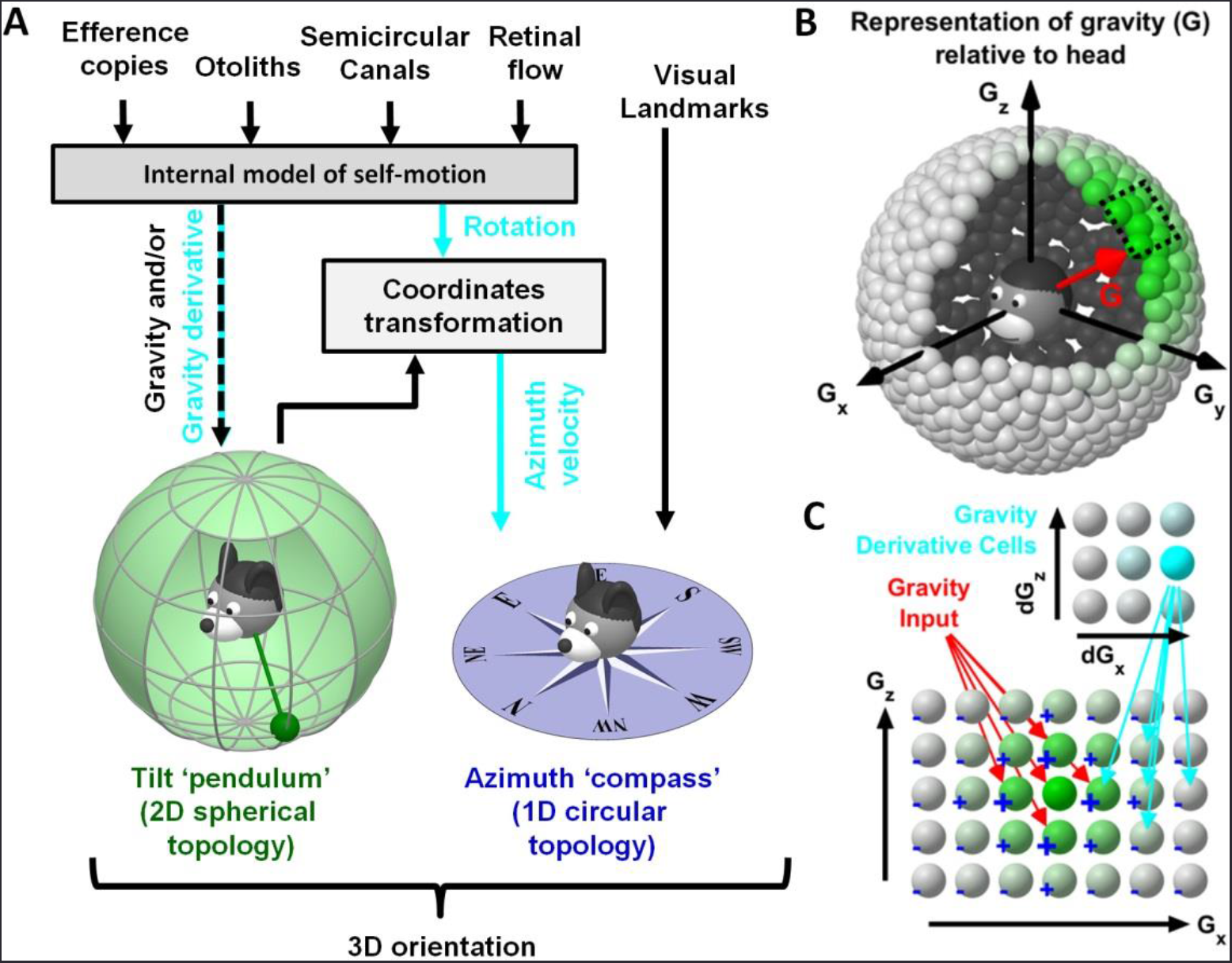
Conceptual model of 3D head orientation compass. **(A)** 3D orientation may be decomposed into 2D tilt and 1D azimuth compasses. Tilt signals are symbolized by a head-fixed pendulum (green). The end of this pendulum may reach any position on a sphere drawn around the head, with each position representing one possible head tilt (spherical topology). Self-motion signals (gravity and/or gravity derivative; and azimuth velocity, cyan) are computed by an internal model of self-motion (Fig. 1A). Note that egocentric rotation signals must be further transformed by gravity to compute allocentric azimuth velocity, necessary to update the ring attractor during 3D rotations (tilted azimuth model). This is because the velocity input to the azimuth compass (HD ring attractor) must include <u>both</u> head-horizontal (yaw, Fig. 6A, cyan) and earth-horizontal (Fig. 6A, green) components. **(B), (C)** Model of 2D attractor that computes 2D tilt. Spherical topology of the proposed model (B), with a portion of the model expanded to show connectivity within the network (C). Cells receive gravity signals (red) and activate neighbors while inhibiting distant cells (’+’ and ‘-’). A gravity-derivative cell tuned to dGx>0 activates rightward connections between gravity-tuned cells, causing the activity to shift. Note the common principles with the 1D model in Fig. 1B. Whether the computation of tilt orientation uses an attractor network remains to be tested experimentally.

An important question for future research is whether tilt signals are processed by a neuronal attractor, similar to azimuth velocity cues. A neuronal attractor for tilt would have a two-dimensional spherical geometry (Fig. 7B), and could integrate inputs that encode the time derivative of gravity in order to track gravity (Fig. 7C). Both of these signals have been identified in the macaque anterior thalamus (Laurens et al., 2016). Yet, it remains unknown whether gravity signals are computed elsewhere and relayed to the anterior thalamus, or computed within a circuit that includes the anterior thalamus itself.

In summary, the importance of gravity on earth life, the existence of 3D HD cells in flying mammals, and the presence of gravity-referenced signals in the macaque (Laurens et al., 2016) and rodent (Cham et al., in preparation) anterior thalamus support the hypothesis that vertical orientation is defined by gravity, computed by or downstream of the brainstem/cerebellar circuit that processes gravito-inertial acceleration (Laurens et al., 2013a,b). HD cells may carry only tilt, only azimuth, or both signals (Finkelstein et al., 2015; Cham et al., in preparation). How these signals interact in individual cells remains to be determined in future experiments.

## NEURAL PATHWAYS

There have been many reviews (Clark and Taube, 2012; Cullen and Taube, 2017; Shinder and Taube 2010; Taube 2007) of the potential pathways that could carry vestibular signals from the medial VN to the lateral mammillary (LMN) and dorsal tegmental nuclei (DTN). Candidate areas carrying vestibular signals to DTN include the nucleus prepositus hypoglossal and the supragenual nucleus (Biazoli et al., 2006; Brown et al., 2005; Butler and Taube, 2015). However, it is possible that vestibular projections influencing HD computation may also arise from the rostral fastigial nuclei, which respond to vestibular stimuli as strongly as the VN (Angelaki et al., 2004; Brooks and Cullen, 2009; Brooks et al. 2015; Shaikh et al., 2005). Projections to DTN and LMN can also arise from reticular nuclei, such as the mesencephalic reticular nucleus, the paragigantocellular reticular nucleus and the gigantocellular reticular nucleus (Brown et al., 2005), which project to the nucleus prepositus hypoglossal (Ohtake 1992) and supragenual nucleus (Biazoli et al., 2006) (reviewed in Shinder and Taube 2010). In fact, fastigiofugal fibers from the rostral part of the fastigial nuclei innervate heavily the nucleus gigantocellularis (NRG) and other reticular nuclei (Homma et al., 1995). Neurons in NRG respond to sensory stimuli of multiple modalities, including vestibular (Martin et al., 2010). Furthermore, electrical stimulation of the rostral NRG evokes ipsilateral horizontal head rotations (Quessy and Freedman, 2004). Preliminary recordings from the dorsal paragigantocellular reticular formation reported neurons modulating similarly during active and passive yaw rotations (Wu Zhou, personal communication), possibly reflecting the total self-motion signal of Fig. 2. Future studies must characterize neuronal properties in these areas, particularly whether they relate to the gravity-dependent properties of the velocity drive to the HD network. In addition, further physiological studies should elucidate whether recurrent activity in the HD networks is modulated in a context-dependent manner.

## HEAD DIRECTION SYSTEM IN INSECTS

Interestingly, insect species possess a HD system, which is located in the ellipsoid body and protocerebral bridge of the central complex, and which bears a striking similarity with the mammalian HD system (Seelig and Jayaraman, 2015). The ellipsoid body is a ring-link structure in which neighboring neurons fire together, forming a ‘bump’ of activity whose position on the ring varies as a function of head direction. Similar to mammalian HD cells, the position of the bump is anchored to visual landmarks when available, and can also track head direction when flies walk in darkness. Furthermore, a recent series of calcium imaging and optogenetic studies (Kim et al., 2017; Green et al., 2017; Turner-Evans et al., 2017) has demonstrated that neuronal activity within the ellipsoid body has the properties of an attractor network, and have identified a neuronal mechanism whereby the protocerebral bridge may provide rotation velocity signals to the ellipsoid body.

The similarity between the neuronal mechanisms that encode head direction in mammals and insects, and the fact that flies sense gravity (Kamikouchi et al. 2009), raise the exciting possibility that some of the computational principles discussed in this perspective may apply to the HD system of insects.

## CONCLUSIONS

We have proposed a quantitative framework for how vestibular signals, and other self-motion cues, contribute to a multisensory estimate of allocentric self-motion, which we hypothesize updates the HD ring attractor (and possibly influences grid cell tuning too). This azimuth velocity signal is influenced by efference copies, canal, visual, somatosensory, as well as gravity signals, thus it requires extensive processing before it can be integrated by the ring attractor. A big challenge in future years, and the only way to quantitatively test the proposed framework, is to experimentally identify the source, neural correlates and properties of this multisensory self-motion estimate. Virtual Reality setups, which provide the ability to independently manipulate visual, vestibular and efference motor/somatosensory signals can be particularly helpful in this regard.

